# Physical activity increases neuronal activity in the circadian clock of diurnal *Arvicanthis ansorgei*

**DOI:** 10.1101/2022.05.31.493966

**Authors:** Rosanna Caputo, Robin A. Schoonderwoerd, Ashna Ramkisoensing, Jan A.M. Janse, Hester C. van Diepen, Sylvie Raison, Paul Pévet, Nienke A.V. Derks, Dominique Sage-Ciocca, Tom Deboer, Etienne Challet, Johanna H. Meijer

**Affiliations:** Department of Cell and Chemical Biology, Laboratory for Neurophysiology, Leiden University Medical Center, P.O. Box 9600, 2300 Leiden, The Netherlands; Institute of Cellular and Integrative Neurosciences, CNRS, University of Strasbourg, 67000 Strasbourg, France; Chronobiotron, UAR3415, CNRS, University of Strasbourg, France

**Keywords:** Diurnality, suprachiasmatic nucleus, *in vivo* electrophysiology, Sudanian grass rat, exercise

## Abstract

2.

The central circadian clock, located in the suprachiasmatic nucleus (SCN) within the brain, regulates daily patterns of activity and physiology. Many studies indicate that exercise at specific times throughout the day can help maintain proper circadian rhythms. In nocturnal animals, even moderate levels of physical activity suppress the neuronal discharge rate of the SCN. Given that such a mechanism would likely be counter-effective in diurnal animals, we measured the firing rate of SCN neurons in freely moving diurnal *Arvicanthis ansorgei* using implanted microelectrodes. We found that SCN firing was acutely increased rather than decreased both during brief (seconds) and long (hours) bouts of activity, and returned to baseline levels after behavioral activity ceased. We also found that daytime activity increases the strength of the SCN rhythm, as expected for day-active animals. To determine whether the acute increases in firing are produced within the SCN or in response to input from outside the SCN, we performed *ex vivo* recordings in which afferent inputs are severed. We found no intrinsic increment occurring in the isolated SCN. These findings suggest that the excitatory effect on the SCN’s neuronal firing rate comes from areas that lie outside the SCN, presumably those that are affected by the animal’s activity. We conclude that exercise has opposite effects on the clock between nocturnal and diurnal rodents, and identified how exercise strengthens the neuronal discharge rhythm in the clock of a diurnal animal.

**Significance statement:** Our biological clock controls behavioral activity rhythms by generating a 24-pattern of electrical activity. The electrical activity serves as output of the clock and is high during the day and low during the night. Physical activity, being under strong control of the clock, acts vice versa, and affects the electrical activity of the clock. In nocturnal animals, behavioral activity inhibits the clock’s firing rate. Here, we examined the effect of behavioral activity on the brain’s clock in the diurnal rodent, *Arvicanthis*. When the animal is active, the clock’s electrical activity is enhanced, rather than decreased. Thus, a diurnal animal can increase the strength of its own clock, by being active during the day.

**Preprint Servers:** The manuscript was deposited as a preprint in bioRxiv preprint doi: https://doi.org/10.1101/2022.05.31.493966; this version posted June 1, 2022. The copyright holder for this preprintin bioRxiv, made available under aCC-BY-NC-ND 4.0 International license (which was not certified by peer review) is the author/funder, who has granted bioRxiv a license to display the preprint in perpetuity. It is made available under aCC-BY-NC-ND 4.0 International license.

**Classification:** Biological Sciences, Physiology

## Introduction

Nearly all life on Earth developed an internal clock that can entrain the timing of physiological processes and behaviors to daily changes in the environment ^1, 2^. In mammals, the timing of physical activity is regulated by the central circadian pacemaker known as the suprachiasmatic nucleus (SCN), a bilateral structure located in the hypothalamus. The SCN generates a rhythm with a period of approximately 24 hours and then conveys this rhythm to the brain and the periphery, thereby establishing synchronized rhythmicity at the systemic level ^1, 3, 4^. Several pathways that convey this functional output from the SCN have been identified and include the autonomous nervous system, circulating levels of glucocorticoids and melatonin, and the timing of behavioral activity ^5^.

Interestingly, in addition to driving clock output, behavioral activity in turn affects our central timing^6, 7^. For example, in humans exercise can induce a phase shift in the circadian clock’s rhythm ^8, 9^, facilitate entrainment to a non-24-hour cycles ^10^, and accelerate realignment to a shifted light-dark cycle ^11, 12^. In nocturnal animals, exercise also induce a phase shift in the circadian system and has an acute suppressing effect on the SCN discharge rate ^13-16^. However, the effect of physical activity on SCN firing has not been examined in diurnal species.

To measure the effects of behavioral input on the SCN in diurnal species, we used the Sudanian grass rat, *Arvicanthis ansorgei*, a diurnal rodent that lives in savannas and grasslands in the Sudano-Guinean region of Africa. In the wild, some animals display crepuscular activity, with increased activity levels at both the beginning and end of the day, while other animal’s activity is distributed throughout the day ^17, 18^. In *Arvicanthis*, the daily rhythm of various parameters such as the plasma levels of glucose, or the levels of the neurotransmitter serotonin, are opposite to nocturnal rodents ^19, 20^. Because *Arvicanthis* belongs to the Muridae family like rats and mice and has a physiology similar to these classic nocturnal rodents, it is an excellent diurnal animal model for comparative studies.

Here, we recorded electrical activity in the SCN of freely moving *Arvicanthis* using implanted electrodes. In addition, we recorded electrical activity in *ex vivo* SCN slices in order to measure the SCN’s intrinsic rhythm in the absence of behavioral input. We found that, unlike nocturnal species, the SCN in diurnal *Arvicanthis* is excited by behavioral activity. Our *ex vivo* recordings of SCN slices revealed a circadian rhythm free from acute changes in firing rate. This indicates that the sudden increments in SCN activity that we measured, do not originate from within the SCN, but are imposed by extra-SCN areas. Importantly, the SCN rhythm is strengthened by physical activity provided it is during the day.

## Results

### The SCN’s multiunit activity waveform reflects the animal’s pattern of behavioral activity

We first recorded multiunit activity (MUA) in the SCN of freely moving *Arvicanthis* (n=11) while simultaneously recording the animals’ behavioral activity. We found that MUA in the SCN was higher during the day than during the night (Supplemental Figure 1). We then determined the shape of the 24-hour rhythm by fitting the curve to all data points in the recordings (Supplemental Figure 2). We found that some recordings had a unimodal MUA rhythm (Figure 1A), while other recordings had bimodal MUA rhythm that was largely bimodal (Figure 1B). In both cases, the MUA rhythm correlated with the animal’s behavioral activity rhythm. We then quantified the bimodality of the MUA and the behavioral activity rhythms in order to obtain a crepuscularity index ranging from 0 to 1, corresponding to the lowest and highest degree of crepuscularity, respectively (Figure 1C). Plotting MUA crepuscularity index against the corresponding behavioral crepuscularity index measured for each animal revealed a significant correlation between the two (Figure 1D). The location of the recording electrode was verified by histological analysis (Figure 1E).

**Figure 1:**
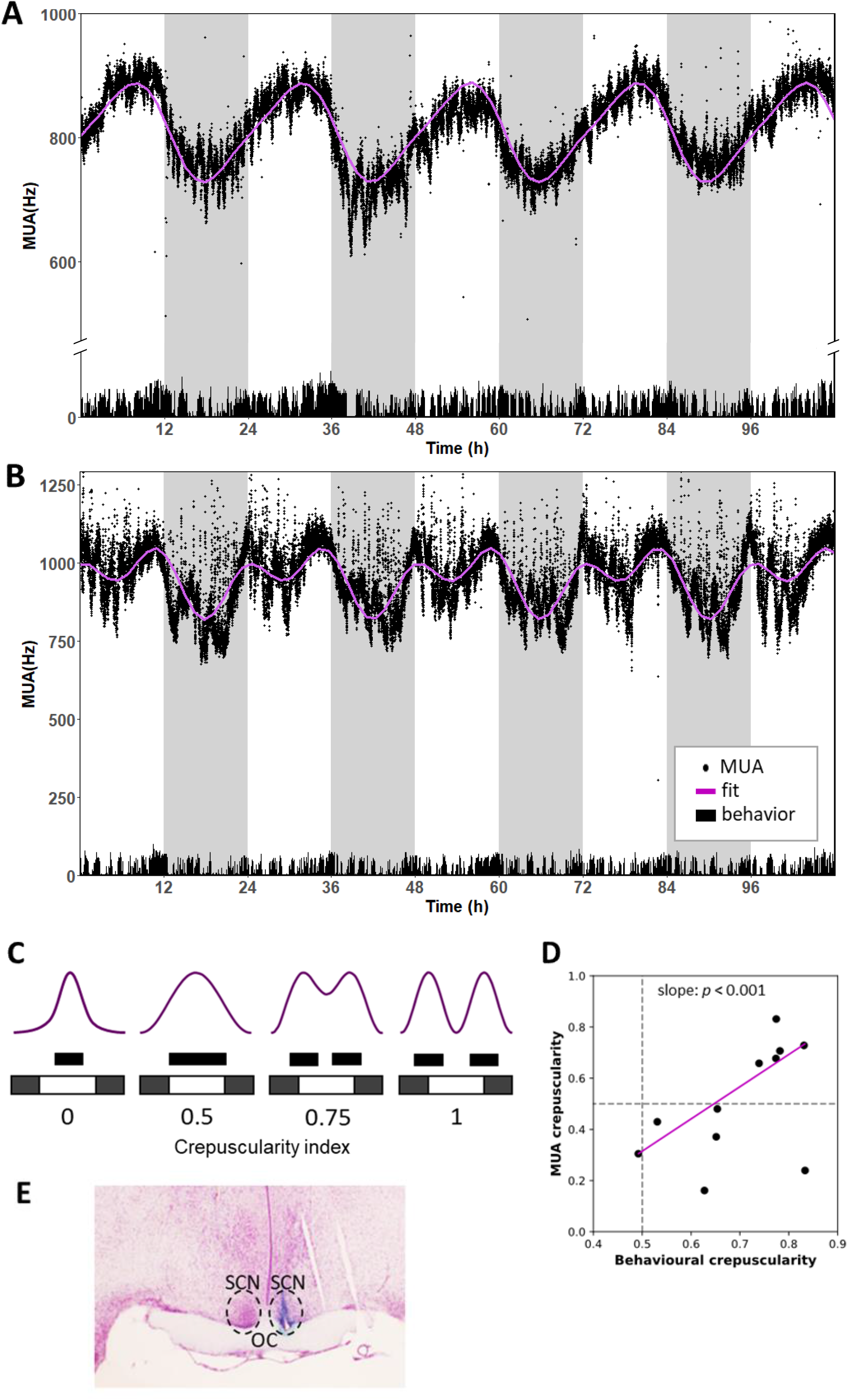
Diurnal *Arvicanthis* display both unimodal and bimodal patterns of SCN electrical activity measured *in vivo*. (A, B) Representative recordings showing an animal with a unimodal multiunit activity (MUA) rhythm (A) and an animal with a bimodal MUA rhythm (B). The solid purple lines show the curves fitted to the datapoints. (C) Schematic depiction of MUA shape (purple line) corresponding to behavioral activity (black blocks) has crepuscularity index ranging from 0 (lowest level of crepuscularity, i.e. no activity in the 6-hour windows surrounding the light-dark and dark-light transitions) to 1 (highest level of crepuscularity). (D) The MUA crepuscularity index was plotted against the behavioral crepuscularity index for each animal, and the linear regression was measured using the Huber loss function. (E) Example of a coronal section of the *Arvicanthis* SCN, located above the optic chiasm (OC). The site of implantation is visualized by the blue spot.

**Figure 2:**
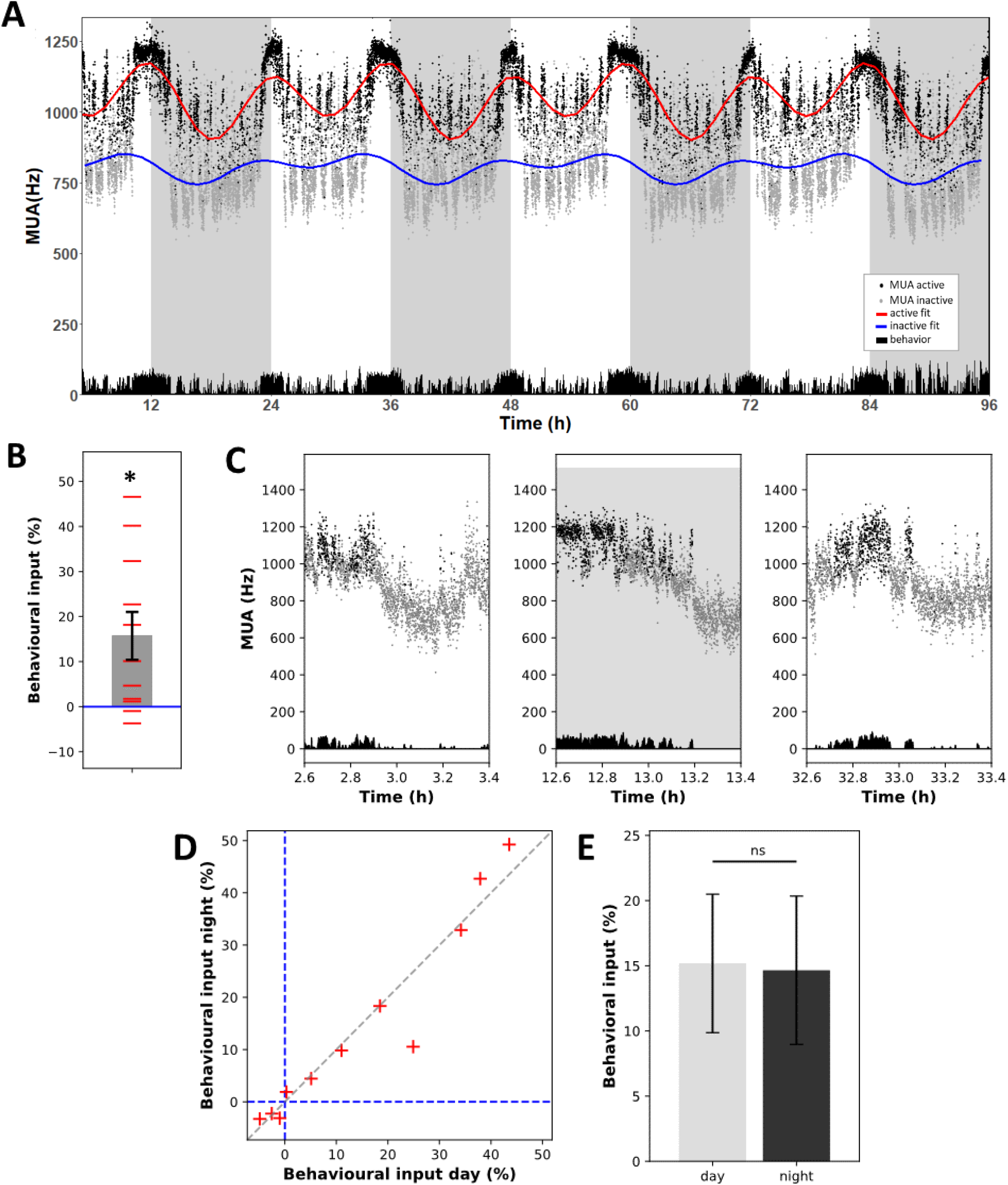
Behavioral activity increases SCN electrical activity measured *in vivo*. (A) Representative recording of multiunit activity (MUA). MUA recorded when the animal is active (MUA active) is represented by black symbols, MUA recorded when the animal is inactive (MUA inactive) is represented by light gray symbols. The red and blue curves were fitted to the active and inactive MUA points, respectively. The gray shaded background indicates darkness, and the animal’s physical activity is indicated in black at the bottom. (B) Each horizontal red line represents the magnitude (expressed as daily mean) of the behavioral input for a given animal. Behavioral input (mean ± SEM), deviates significantly (one-sample T-test, *p≤0.05) from 0 (blue line). (C) Representative magnified time courses of the MUA recording showed in figure A, measured during bouts of physical activity.(D) The magnitude of behavioral input during the night plotted against the magnitude of behavioral input measured during the day for each animal; the dashed gray line represents the 1:1 correlation (*y*=*x*). (E) Summary (mean ± SEM) of the magnitude of behavioral input measured during the day and during the night (n=11 animals); ns, not significant.

### Behavioral activity increases the electrical activity of SCN neurons

Next, we analyzed the MUA data separately when the animal was active and when the animal was not active. On an ultradian (i.e., minutes-hours) timescale, upward deflections were observed at the times in which the animal was active (Figure 2A). We then fit a curve to all timepoints measured when the animal was inactive, revealing a daily rhythmic profile. The curve fitted to all active timepoints showed a rhythmic profile as well, but at a higher level (Figure 2A); this is because on average, MUA was higher at the timepoints at which the animals were active (Figure 2B). At the onset of physical activity, we observed an acute increase in SCN neuronal firing, while we observed a gradual decrease in SCN firing rate at the offset of physical activity (Figure 2C). The increases in firing rate were present both during the day and night (Figure 2D), and the magnitude was similar between day and night (Figure 2E).

### The intrinsic SCN rhythm is devoid of higher frequencies

Next, to determine whether the acute increases in MUA frequency are generated within the SCN we prepared *ex vivo Arvicanthis* brain slices containing the SCN but lacking neuronal input from other brain structures. We recorded slices prepared from 8 animals that displayed both unimodal and crepuscular behavioral activity. In none of the *ex vivo* recordings sudden transitions in MUA levels were observed (Figure 3) such as those recorded in vivo, but instead, smooth sinusoidal rhythms were observed in all our *ex vivo* recordings. Thus, the sudden transitions are not produced in the isolated SCN.

**Figure 3:**
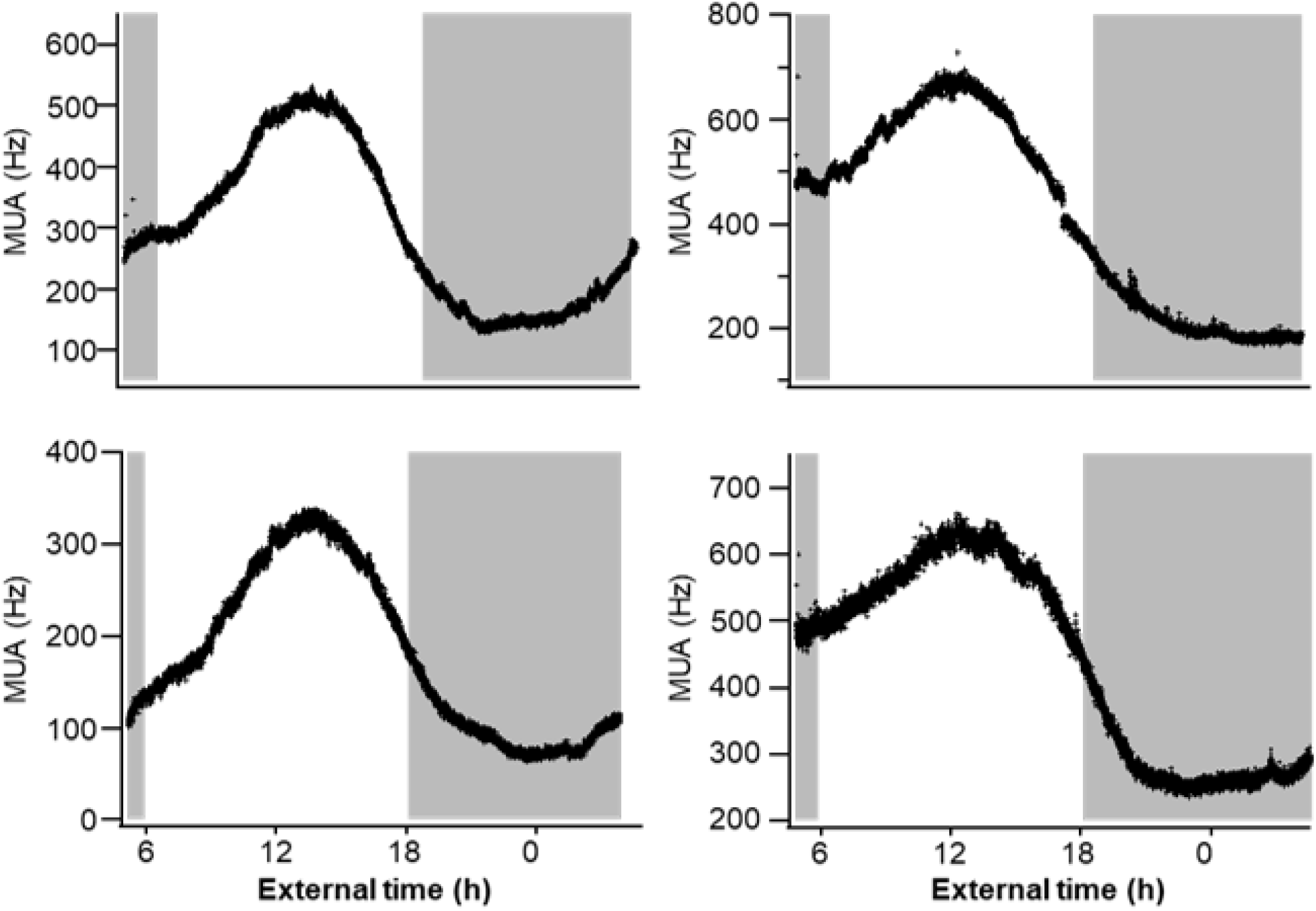
Intrinsic SCN electrical activity is devoid of higher frequencies. Representative recordings of *ex vivo* brain slices containing the *Arvicanthis* SCN. The shaded gray areas indicate the periods of darkness to which the animals were entrained prior to preparing the *ex vivo* slices.

### Behavioral input increases the amplitude of the SCN rhythm

Given the excitatory effects of behavioral activity on MUA measured in the SCN of *Arvicanthis*, we hypothesized that behavioral input would affect the amplitude of MUA rhythms in the SCN of diurnal species. Thus, in diurnal animals activity during the day would likely increase MUA activity to higher levels, while inactivity during the night would keep MUA at low levels (shown schematically in Figure 4A, example provided in Figure 4B). In *Arvicanthis* showing a crepuscular behavioral activity, MUA levels would be particularly increased at the beginning and at the end of the day, resulting in a bimodal MUA waveform (shown schematically in Figure 4C, example provided in Figure 4D).

**Figure 4:**
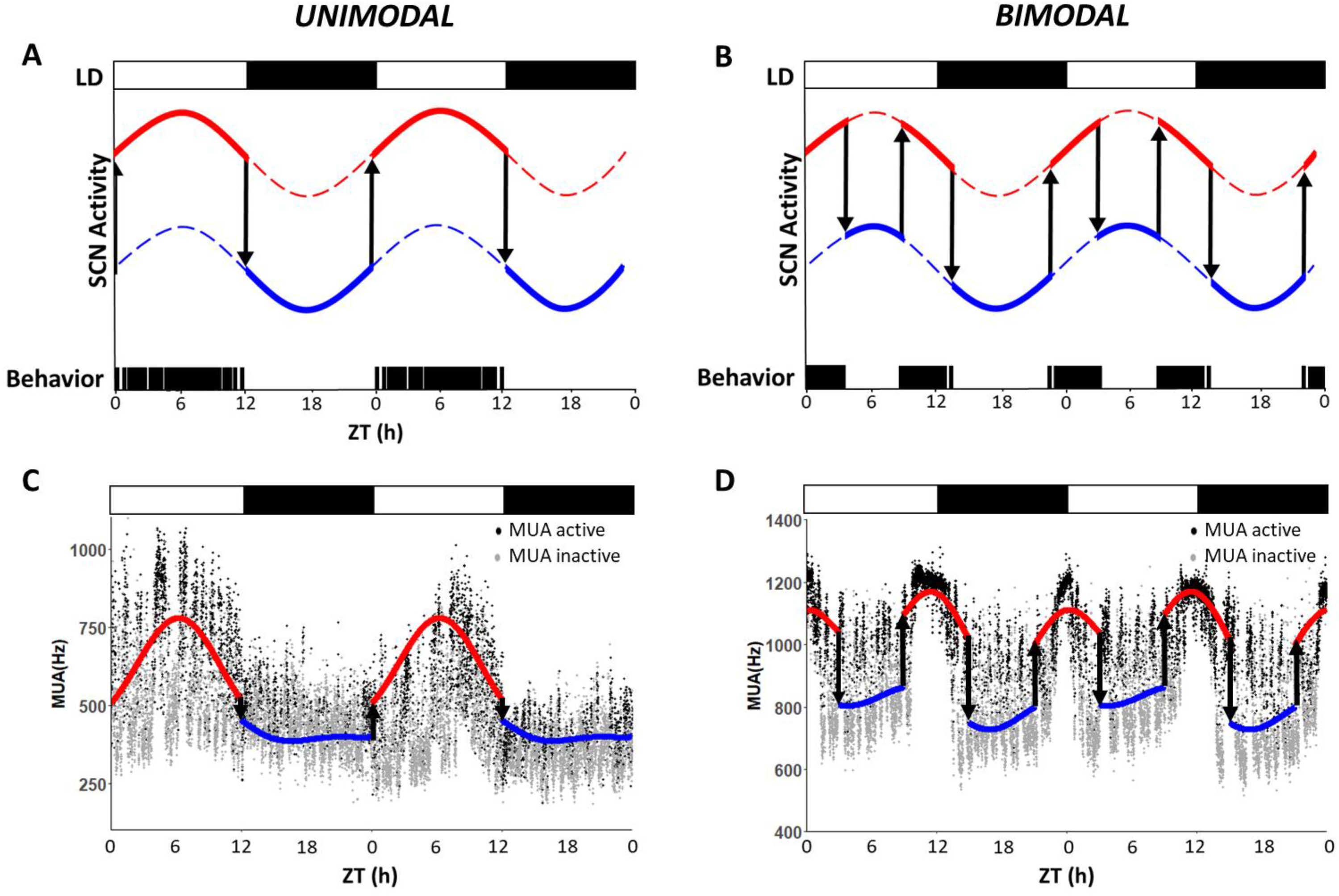
Schematic depiction of unimodal and bimodal SCN MUA pattern. Interpolating lines of SCN MUA during periods of activity (red line) and periods of inactivity (blue line), in a schematic depiction of a unimodal (A) and bimodal (B) SCN MUA pattern. (A) Activity during the day, indicated by the black bars at the bottom of the graph, increases the amplitude of the total MUA rhythm (red and blue bold solid lines). (B) Activity during the dawn and dusk, indicated by the black bars at the bottom of the graph, increases MUA levels and confer a bimodal shape to the SCN MUA rhythm. (C) To visualize the effect of a unimodal behavioral activity on a SCN MUA recording, the fitted line (red line) of MUA during activity (black points) is plotted during the day and the fitted line (blue line) of MUA during inactivity (gray points) is plotted during the night. (D) To visualize the effect of a crepuscular behavioral activity on a SCN MUA recording, the fitted line (red line) of MUA during activity (black points) is plotted during the light/dark and dark/light transitions and the fitted line (blue line) of MUA during inactivity (gray points) is plotted in the middle of the day and of the night. The light-dark cycle (LD) is shown above the graphs. ZT, Zeitgeber time, with ZT0 and ZT12 corresponding to lights on and lights off, respectively.

## Discussion

Although exercise has been shown to inhibit SCN neuronal activity in nocturnal species such as hamsters ^13^, rats ^14, 15^, and mice ^16^, the effect of activity on the SCN in diurnal animals has not been investigated ^7^. To address this question, we measured the effect of behavioral activity on the *in vivo* firing rate of SCN neurons in freely moving diurnal *Arvicanthis ansorgei*. Unlike in nocturnal species, physical activity increases neuronal activity in the SCN of *Arvicanthis*. This effect seemed superimposed on the SCN’s intrinsic circadian rhythm in discharge rate, as it was absent *ex vivo*, in acutely prepared hypothalamic brain slice. In unimodal diurnal animals, the elevating effect of behavioral activity resulted in a higher peak in SCN activity at midday; in crepuscular *Arvicanthis*, the result was an enhanced bimodal pattern of SCN electrical activity. Based on these results, we conclude that: *i*) behavioral activity excites the SCN of diurnal animals *ii*) the effect of behavior is opposite in diurnal compared to nocturnal animals, and *iii*) behavioral activity during the day is effective in enhancing the SCN amplitude, by raising the peak in electrical activity that occurs during the day. Since the electrical activity of the SCN functions as clock’s output, physical activity can strengthen our internal clock.

Both in the wild and in laboratory conditions, the physical activity of some *Arvicanthis* show a crepuscular pattern, with higher activity levels during the beginning and the end of the day ^17, 18^. A crepuscular pattern of behavioral activity has been described in many diurnal rodents, such as the Octodon degus ^21^ and the Mongolian gerbils ^22^, but also in other diurnal species, such as in the Drosophila^23^. Despite the crepuscular pattern of activity of some individuals, *Arvicanthis* physiology displays features typical for diurnal species. First, *Arvicanthis* retinae consist of a high percentage (∼33%) of cone photoreceptors^24^, in contrast to less than 3% in mice^25^ and rats^26^. Cones are indeed fundamental for diurnal species depending much on daytime vision and color detection. Morover, the circadian rhythm in arousal structures like the serotonergic raphe nuclei or the noradrenergic locus coeruleus is opposite between *Arvicanthis* and nocturnal rats^20, 27, 28^.

In all mammals measured so far, the firing rate of SCN neurons is high during the day and low during the night based on *in vivo* recordings in nocturnal rodents such as hamsters ^13^, rats ^14, 15, 29^, and mice ^16^, as well as in diurnal chipmunks ^30^. These findings have been confirmed using *ex vivo* SCN recordings obtained from several nocturnal rodents ^14, 31, 32^ and diurnal *Rhabdomys* ^33^. Now we confirm this pattern in diurnal *Arvicanthis*.

The incremental effects of physical activity on MUA measured in diurnal *Arvicanthis* (up to 46.6%) are similar in magnitude to the inhibition reported for nocturnal species (ranging from 5-60%) ^15^. We found that 6 out of 11 *in vivo* recordings (54.5%) in our study showed behavioral input (based on an arbitrary threshold of a 10% change at active timepoints relative to inactive timepoints). Also in rats and mice, responses to behavior were not found in all recordings. Most likely, the electrode location is not always in the SCN areas receiving the behavioral feedback. This would resemble the organization of the light input pathway that reaches only the ventral part of the SCN ^34^. As a less likely but possible explanation is that not in all animals the SCN receives feedback from activity. In addition, and in contrast to observations reported in nocturnal species ^15^, the magnitude of the behavioral input that we measured in *Arvicanthis* was similar between day and night. One possible explanation for the difference between nocturnal and diurnal animals may be that in nocturnal species the relatively low firing rate of SCN neurons cannot be decreased further by the inhibitory effects of behavioral activity (i.e., a floor effect), while the firing rate in the SCN of diurnal *Arvicanthis* may not have reached a ceiling level and can be increased further by behavioral input.

For an animal that is typically active during the day and inactive during the night, excitatory behavioral input can increase the amplitude of the SCN’s neuronal activity, provided that the animal is active during the day and inactive during the night. The increment in amplitude of the SCN rhythm by behavior also occurs in nocturnal species, but with the opposite mechanism. In nocturnal animals, inhibitory inputs cause an increase in amplitude by lowering the trough of the SCN rhythm. As discussed above, all nocturnal species studied to date have a 24-hour sinewave rhythm in SCN electrical activity measured *in vivo* ^13-16, 29^, whereas many of our *in vivo* recordings in *Arvicanthis* had a bimodal rhythm in the SCN. This can be explained by the animals’ crepuscular behavioral activity (increased physical activity during dawn and dusk).

Our data show that the acute increases in firing rate observed in the *in vivo* recordings are not present in *ex vivo* recordings. This suggests that behavioral activity increases SCN firing rate rather than the SCN stimulates acute activity. A limitation of our study is that we cannot exclude the possibility of a brain area that stimulates both electrical and behavioral activities. The physical activity is a major feedback to the SCN, but it is not the only feedback. Indeed, other behavioral activities (such as eating, grooming, etc.) feedback to the SCN as well and induce changes in SCN MUA levels, albeit to a lesser extent (see Oosterhout et al., 2012^16^). The only case where there is a complete absence of any input is in slices recorded *ex vivo* ^14^. When we recorded the *Arvicanthis* SCN *ex vivo*, the recording traces were devoid of any acute increase.

A remarkable feature of behavioral input is that the MUA increased sharply at the start of the behavior onset, but returned to baseline slowly after activity ceased. The same kinetics, albeit in opposite direction, has also been observed in nocturnal rodents ^15, 16^ and likely reflects the dynamics of a relatively slow-acting G protein–coupled receptor such as a neuropeptide receptor ^35^. The most likely candidates for mediating this effect are neuropeptide Y (NPY) released from the intergeniculate leaflets of the thalamus (IGL) ^3, 36^ and serotonin released from the raphe nuclei ^37^, as both IGL lesions^38^ and ablation of serotonergic afferent SCN pathways ^39, 40^ have been shown to reduce behavior-induced phase shifts. We therefore hypothesize that NPY and serotonin also mediate the effects of behavioral input in diurnal species, but have an opposite effect on MUA compared to nocturnal species. Injections of serotonin receptor agonists in day-active *Arvicanthis* induce a small phase advance during the subjective night ^41^, while in nocturnal rodents this phase advance occurs during the subjective midday ^42^. The relationship between NPY and behavioral input is largely unknown in diurnal species; however, NPY-containing neurons in the IGL, which project to the SCN, are activated by wheel running ^43^.

Because behavioral input excites the SCN in diurnal species, the phase-response curve of exercise is expected to be similar to the phase-response curve induced by light pulses ^7, 44^. Indeed, in humans exercise advances our rhythm when performed in the early morning and is delaying in the late evening and early night ^9, 45^. We therefore speculate that in diurnal species, physical activity and light act in synergy, as both inputs lead to the same phase response curve and both light and behavior have excitatory effects on the SCN ^46^ (Table 1). This is in contrast to nocturnal animals where light and behavioral activity have opposing effects on the SCN (Table 1) and on phase shifts ^15^.

**Table 1:** Effect of light and behavior on the SCN in diurnal and in nocturnal species.

|  | Diurnal |  | Nocturnal |  |
| --- | --- | --- | --- | --- |
|  | + | - | + | - |
| *Light* | x | x | x |  |
| *Behavior* | x |  |  | x |

In diurnal animals, the SCN is both excited (+) and inhibited (-) by light, and is excited by behavior. In contrast, in nocturnal animals, the SCN is excited by light and inhibited by behavior. Light and behavior work in synchrony in diurnal, but antagonistically in nocturnal species.

In summary, we report that the SCN in diurnal *Arvicanthis ansorgei* responds acutely to behavioral input with increased firing that is sustained for the entire duration of the physical activity, thereby raising the electrical activity to higher levels during daytime and increasing the amplitude of SCN firing. Consequently, the SCN rhythm and the behavioral activity rhythm form a positive feedback loop. Finally, given the positive effects of properly timed exercise on the circadian system in humans, future studies are warranted in order to identify the neuronal pathway by which behavioral information feeds back to the SCN.

## Materials and Methods

### Animals

Adult (3-6-month-old) Sudanian grass rats (*Arvicanthis ansorgei*) of both sexes were obtained from Chronobiotron UMS 3415 (Centre National de la Recherche Scientifique, University of Strasbourg, France) and used for *in vivo* (n=39 animals) and *ex vivo* (n=8 animals) experiments. The animals were housed in transparent plastic cages under a 12:12-hour light-dark (LD) cycle consisting of 200 lux white light <5 lux red dim light (darkness); food and water were provided *ad libitum*. Locomotor activity was recorded using a passive infrared (PIR) sensor. All experiments were performed in accordance with the guidelines established by European Commission Directive 2010/63/EU and the French Ministry of Higher Education and Research (APAFIS #6099-2016090216536741, v4).

### In vivo electrophysiology

Multiunit activity (MUA) was recorded in the SCN of freely moving animals as described previously ^47^. Tripolar stainless steel microelectrodes (model MS333-3-BIU-SPC; Plastics One, Roanoke, VA) were implanted in the SCN of animals anesthetized with Rompun (4 mg/kg body weight; Bayer Pharma, Puteaux, France). In brief, two polyamide-insulated twisted electrodes (bare electrode diameter: 125 µm) for differential recordings were aimed at the SCN using a stereotactic frame with an angle of 5° in the coronal plane, with the following coordinates: 0.7 mm anterior to Bregma, 0.9 mm lateral to the midline, and 7.8 mm ventral to the dura mater. A third uncoated electrode was placed in the cortex as a reference. The electrodes were fixed to the skull using screws and dental cement. After at least 5 days of recovery, the animals were connected to a custom-designed recording setup in which SCN extracellular activity was recorded while the animal was able to move freely. The electrical signal was amplified and bandwidth filtered (500-5000 Hz), and window discriminators were used to convert action potentials to digital pulses that were counted in 10-second or 1-second bins using custom-made software. The differential amplifier was based on the design reported by Yamazaki and colleagues ^13^. The recording chamber was also equipped with a PIR sensor to measure behavioral activity. Behavioral data were recorded in exactly the same 10-second or 1-second bins as the MUA activity, using an integrated recording setup.

At the end of the experiment, the location of the microelectrode in the SCN was verified by histology. In brief, the animal was euthanized with an overdose of sodium pentobarbital (3 ml/kg body weight; CEVA, Libourna, France) under inhalation anesthesia with 4% isoflurane in O_2_. The recording site was marked by delivering a small electrolytic current (40 µA) through the recording electrode to deposit iron at the electrode tip. The brain was then removed and fixed for 2 days in 4% paraformaldehyde containing C_6_FeK_4_N_6_·3H_2_0 (Sigma-Aldrich, Lyon, France). Coronal slices (40 µm thick) were then prepared, and the recording site was reconstructed.

Surgery for electrode implantation was performed on 39 animals; 15 implantations successfully targeted the SCN (success rate of implantation was ≃ 38%). Recording from 11 animals were kept, 4 were discarded because of instability of the signal.

### Ex vivo electrophysiology

MUA was recorded in the SCN as described previously ^48^. The *Arvicanthis (n= 8)* were entrained to a 12:12-hour LD cycle for at least 30 days prior to the experiment. The animals were then sacrificed by decapitation under a dim red light at Zeitgeber time (ZT) 21±0.5 h, which corresponds to three hours before lights on. The brains were removed within one minute of decapitation, and brain slices (∼450 μm thick) were prepared using a tissue chopper. The slice containing the SCN was transferred to a laminar flow chamber within six minutes of decapitation and bathed in oxygenated artificial cerebrospinal fluid (ACSF) containing (in mM): NaCl 116.4, KCl 5.4, NaH_2_PO_4_ 1.0, MgSO_4_0.8, CaCl_2_1.8, NaHCO_3_ 23.8, 15.1 glucose, and 5 mg/L gentamicin (Gibco) saturated with 95% O_2_ and 5% CO_2_ (pH 7.2-7.4 and 290-310 mOsm). The slices were stabilized using an insulated tungsten fork and maintained for 1 hour prior to placing a 90% platinum and 10% iridium electrode (50-µm diameter) in the center of the SCN. The electrical signal was amplified 10x and bandpass-filtered (0.3-3000 Hz). Action potentials that exceeded a predetermined threshold set well above noise (∼5 μV) were counted in 10-second bins using custom-made software.

### Data analysis

Data from the *ex vivo* electrophysiology experiments were analyzed using a custom-made program written in MATLAB. Multiunit recordings lasting at least 24 hours and expressing a clear peak in MUA were moderately smoothed using a least-squares algorithm ^49^.

Data from the *in vivo* electrophysiology experiments were analyzed using Python 3.0.9 with the Pandas module version 1.3.0, and visualized using Matplotlib version 3.4.2, or RStudio version 1.4.1103. A 48-hour window consisting of 2 LD cycles starting with the onset of a light phase was used for further analysis; within this window, the following formula was fitted to the MUA datapoints:

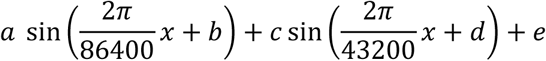

This formula combines a sine curve with a 24-hour period (i.e., 86,400 seconds) to fit the circadian component in the MUA rhythm with a sine curve with a 12-hour period (i.e., 43,200 seconds) in order to fit the crepuscular components of the MUA rhythm. This formula was fitted separately to the timepoints coinciding with behavioral activity (indicated by the red lines in Figures 1 and 4) and to the timepoints coinciding with behavioral inactivity (indicated by blue lines in Figures 1 and 4). The whole recording bin was scored as active if it contained at least 1 movement count of the PIR sensor and scored as inactive if no PIR counts were measured. In the same 48-hour window, the average MUA was calculated for all time points coinciding with behavioral activity and for all time points coinciding with behavioral inactivity. Behavioral input was calculated by dividing the average MUA of timepoints with activity by the average MUA of timepoints with inactivity, expressed as the percent change. To determine whether the magnitude of the behavioral input differed between day and night, the timepoints obtained at both phases of the cycle were analyzed separately.

To analyze crepuscular behavior, we calculated a “crepuscularity index” by dividing the level of activity in the 6-hour windows surrounding the light-dark and dark-light transitions by the total level of activity measured throughout the recordings. The MUA crepuscularity was calculated by fitting the formula described above to all MUA datapoints and then dividing the absolute value of the 12-hour sine curve amplitude *(c)* by the sum of the absolute values of the 24-hour sine curve amplitude (*a*) and *c*, as shown below:

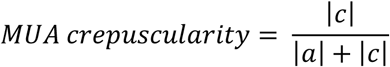

A crepuscularity index of 0 and 1 corresponds to no crepuscularity and complete crepuscularity, respectively.

### Statistics

Data were analyzed using SPSS, Sigma Plot, or the Python module SciPy version 1.7.0. To determine whether the magnitude of behavioral input differed significantly from 0, we used a one-sample Student’s *t*-test. Differences in behavioral input between the day and night were analyzed using a paired Student’s *t*-test. The correlation between the MUA crepuscularity index and the behavioral crepuscularity index was assessed using robust linear regression with the Huber loss function as described previously ^50^. All summary data are presented as the mean ± the standard error of the mean (SEM), and differences were considered significant at *p*≤0.05.

## 5. Acknowledgments and funding sources

We thank Sylviane Gourmelen, René Wilbers, and Maxime Houtekamer for assistance and technical support. This publication is part of the project BioClock (with project number 1292.19.077) of the research programme Dutch Research Agenda: Onderzoek op Routes door Consortia (NWA-ORC) which is (partly) financed by the Dutch Research Council (NWO), obtained by J.H.M. This research was supported by the NWO (Netherlands Organization for Scientific Research; Complexity grant number 645.000.010 to J.H.M.), the Velux Stiftung foundation, Zurich, Switzerland (project number 1131 to J.H.M.), the European Research Council Advanced Grant (project number 834513 to J.H.M.), the initiative of excellence IDEX-Unistra from the French national program “Investment for the future” (ANR-10-IDEX-0002-02 to R.C. and S.R.), and grants from the Centre National de la Recherche Scientifique and the University of Strasbourg (to E.C. and S.R.).

## Author contributions

J.H.M., R.A.S, R.C. designed research; R.C., R.A.S., A.R., H.C.D., N.A.V.D., performed research; J.A.M.J., D.S.C., P.P. contributed new reagents or analytic tools; R.A.S., R.C. analyzed data; R.A.S., J.H.M., R.C. wrote the paper; R.C., R.A.S, A.R., H.C.D., S.R., P.P., N.A.V.D., D.S.C., T.D., E.C., J.H.M reviewed the paper.

## Data Availability Statement

The data that support the findings of this study are available upon request from the corresponding author.

