## Supplementary_data for "Physical activity increases neuronal activity in the circadian clock of diurnal *Arvicanthis ansorgei*"

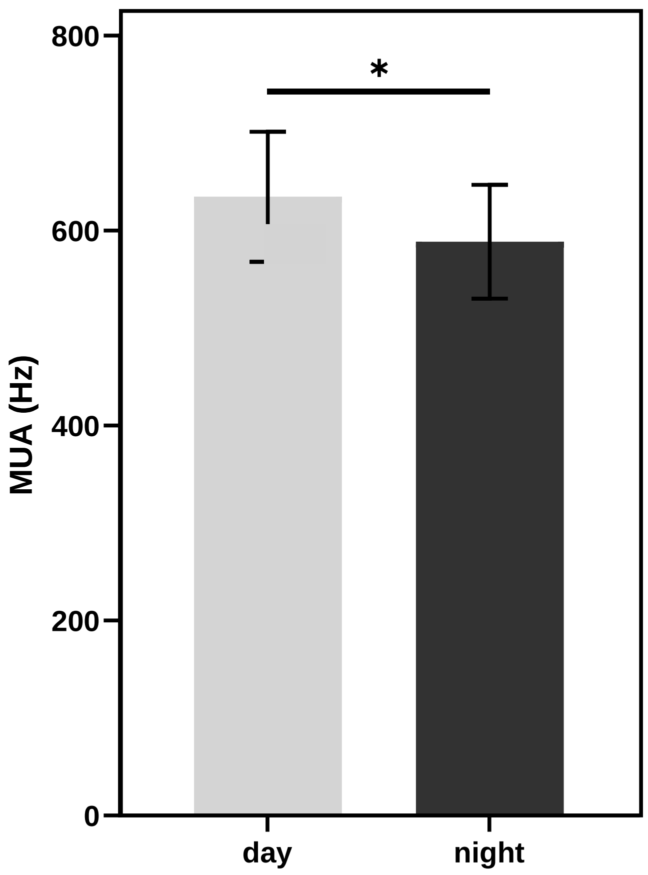


**Supplemental Figure 1: Mean levels of SCN MUA during the day and during the night.** Animals n=11. In *Arvicanthis*, MUA levels during the day are significantly (paired Student's *t*-test *p≤0.05) higher than during the night.

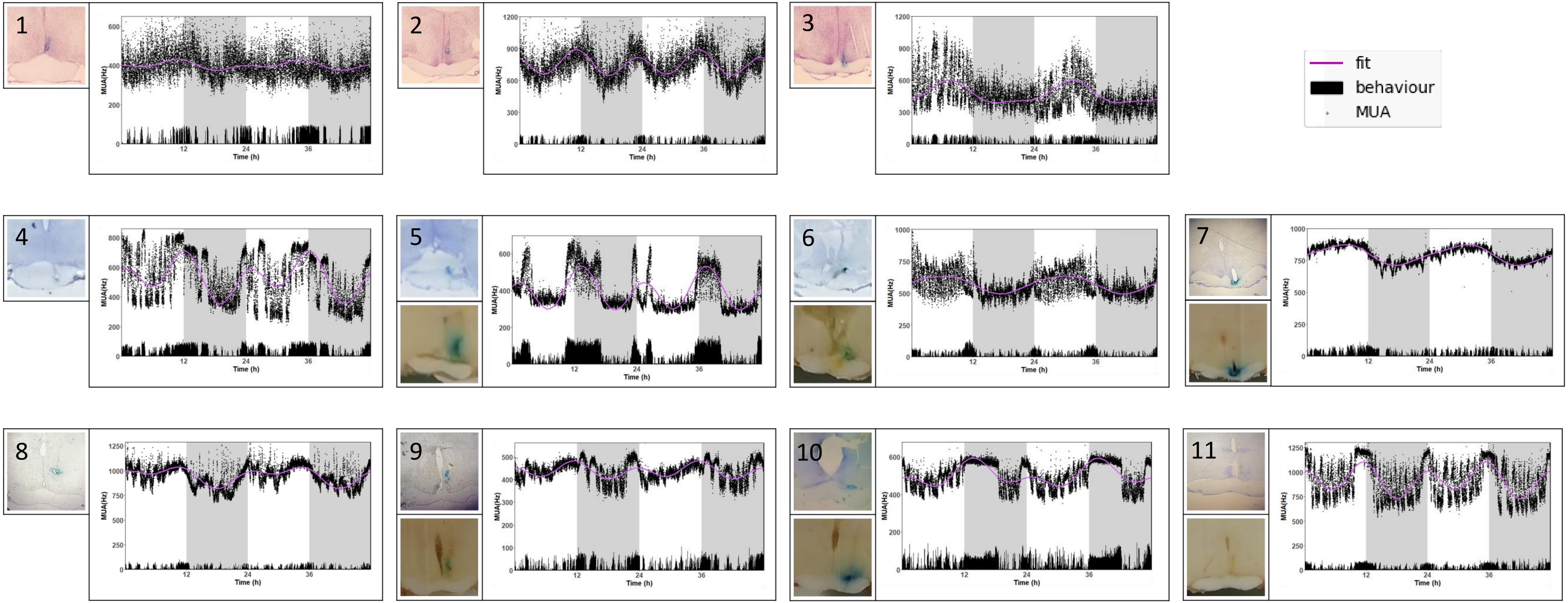


**Supplemental Figure 2: Histology and SCN MUA recording.** Histology of electrode implantation in the SCN of all 11 *Arvicanthis ansorgei,* flanked by the corresponding animal's recording. In each recording MUA (black points) and behavioral activity (black lines) are plotted in 10s bins. The solid purple lines show the curves fitted to the data points.

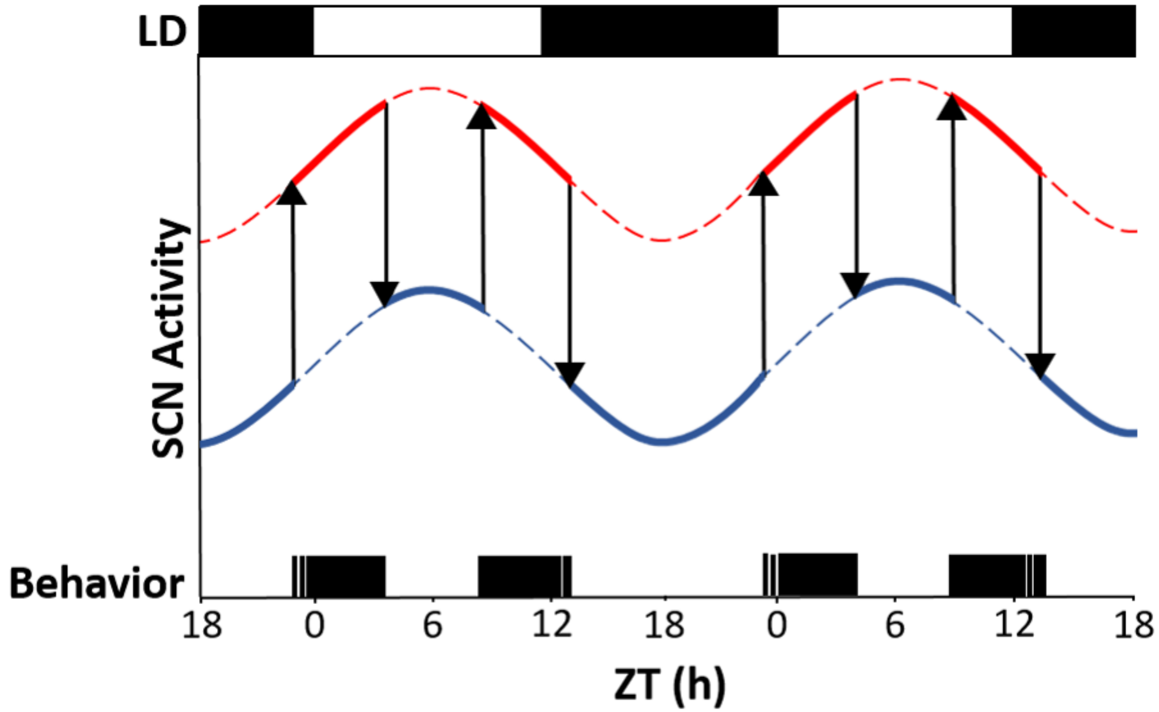


**Supplemental Figure 3: Schematic depiction of SCN MUA pattern in crepuscular animals.** Interpolating lines of SCN MUA during periods of activity (red line) and periods of inactivity (blue line). Activity during the dawn and dusk, indicated by the black bars at the bottom of the graph, increases MUA levels (red and blue bold solid lines) and confer a bimodal shape to the SCN MUA rhythm. The light-dark cycle (LD) is shown above the graph. ZT, Zeitgeber time, with ZT0 and ZT12 corresponding to lights on and lights off, respectively.
